# *In vivo* assessment of mechanical properties during axolotl development and regeneration using confocal Brillouin microscopy

**DOI:** 10.1101/2022.03.01.482501

**Authors:** Camilo Riquelme-Guzmán, Timon Beck, Sandra Edwards-Jorquera, Raimund Schlüßler, Paul Müller, Jochen Guck, Stephanie Möllmert, Tatiana Sandoval-Guzmán

## Abstract

In processes such as development and regeneration, where large cellular and tissue rearrangements occur, cell fate and behavior are strongly influenced by tissue mechanics. While most well-established tools probing mechanical properties require an invasive sample preparation, confocal Brillouin microscopy captures mechanical parameters optically with high resolution in a contact-free and label-free fashion. In this work, we took advantage of this tool and the transparency of the highly regenerative axolotl to probe its mechanical properties *in vivo* for the first time. We mapped the Brillouin frequency shift with high resolution in developing limbs and regenerating digits, the most studied structures in the axolotl. We detected a gradual increase in the cartilage Brillouin frequency shift, suggesting decreasing tissue compressibility during both development and regeneration. Moreover, we were able to correlate such increase with the regeneration stage, which was undetected with fluorescence microscopy imaging. The present work evidences the potential of Brillouin microscopy to unravel the mechanical changes occurring *in vivo* in axolotls, setting the basis to apply this technique in the growing field of epimorphic regeneration.

## INTRODUCTION

Regeneration is the ability to repair and regrow injured or lost body parts, allowing the re-establishment of the missing structure and its functionality. Although an advantageous feature in some multicellular organisms, the potential for regeneration has greatly diverged throughout evolution (Bely and Nyberg, 2010). Among vertebrates, the most remarkable regeneration abilities are reported in urodele amphibians (salamanders and newts) as they are capable of restoring complex structures throughout their lives, such as the retina (Eguchi et al., 2011), heart (Godwin et al., 2017), central nervous system (Albors et al., 2015) and appendages (Iten and Bryant, 1973; Iten and Bryant, 1976; Spallanzani, 1769). Conversely, regeneration abilities in mammals are mainly observed in organs and tissues, and in most cases, only a partial restoration of the structure and function is achieved after injury (Iismaa et al., 2018). However, regeneration of complex structures have been reported in digit tips in mice (Akash et al., 2011) and human children (Illingworth, 1974).

Considering the remarkable regeneration abilities of urodeles, an extensive body of research has focused on understanding the process of limb regeneration and its regulation, with the ultimate goal of translating such knowledge into the development of regenerative therapies in humans (Stocum, 2017). Our knowledge on gene expression and cell transdifferentiation trajectories has advanced considerably in the recent years (Currie et al., 2016; Currie et al., 2019; Gerber et al., 2018; Knapp et al., 2013; Qin et al., 2021; Sandoval-Guzman et al., 2014); however, the influence of the extracellular environment and tissue mechanical properties have remained largely unexplored.

Cells sense and integrate mechanical cues that regulate their fate and behavior (d’Angelo et al., 2019; Wagh et al., 2021). Indeed, mechanical signals direct different stages of embryogenesis (De Belly et al., 2021; Muncie et al., 2020; Nikolopoulou et al., 2017; Weberling and Zernicka-Goetz, 2021), cell fate during organogenesis (Brownfield et al., 2013; Datta et al., 2006; Dietrich et al., 2014; Nelson et al., 2017; Yamamoto et al., 2005) and loss of stemness during aging (Lynch et al., 2017; Segel et al., 2019). Moreover, mechanical properties play key roles in pathological conditions such as fibrosis and cancer (Barnes et al., 2018; Bataller and Brenner, 2005; Northey et al., 2020). Comparatively, little is known about the role of tissue mechanics on animal regeneration. Considering the similarities with embryogenesis, particularly regarding large-scale tissue movements and patterning, it is very likely that physical forces and properties play an equally relevant role (reviewed in Vining & Mooney, 2017). Therefore, complementing our cellular and molecular knowledge with biomechanical measurements will improve our understanding of limb regeneration.

A variety of techniques have been developed to quantify the mechanical properties of biological samples (Akhtar et al., 2011). These methods however, present some limitations that hinder their use for *in vivo* measurements, such as the need for direct contact with the cells of interest or rather poor three-dimensional and/or subcellular resolutions. To overcome such limitations, confocal Brillouin microscopy has emerged as a non-destructive, label-free and contact-free method that can probe viscoelastic properties of biological samples in three dimensions (reviewed in Prevedel et al., 2019). This technique is a type of optical elastography, originally developed in the context of material science, which became widely used for the study of condensed matter (Dil, 1982) and was combined with scanning confocal microscopy in the last decades for the measurement of biological samples (Scarcelli and Yun, 2008). Confocal Brillouin microscopy utilizes the inelastic scattering of light due to interactions between incident photons and spontaneous thermally-induced density fluctuations, which can be described as a population of microscopic acoustic waves, called phonons. The scattering process causes an energy transfer between the phonons and the incident photons, which is detectable as a frequency difference between incident and scattered light (the Brillouin frequency shift). Importantly, the sound-wave properties exhibit an intrinsic dependence on the material’s viscoelastic properties, i.e., the longitudinal modulus and the viscosity. The viscosity of the material will attenuate the propagation of the acoustic wave due to energy dissipation, and is proportional to the linewidth of the Brillouin scattered light spectrum. The longitudinal modulus is proportional to the Brillouin frequency shift and describes a material’s elastic deformability under a distinct type of mechanical loading (i.e., compressibility). The Brillouin frequency shift furthermore depends on the sample’s refractive index and density. Notably, it has been shown that changes in the Brillouin frequency shift typically arise from changes in the longitudinal modulus and that the influence of changes in refractive index and density may be negligible for samples with low lipid content (Scarcelli et al., 2015; Schlüßler et al., 2018; Schlüßler et al., 2022).

Thus far, confocal Brillouin microscopy has been used for the measurements of viscoelastic parameters in the eye and cornea (Scarcelli and Yun, 2012; Vaughan and Randall, 1980), cancer tumors and organoids (Conrad et al., 2019; Margueritat et al., 2019), bone and cartilage (Akilbekova et al., 2018; Alunni Cardinali et al., 2021; Mathieu et al., 2011), neurons and glia (Schlüßler et al., 2018), among other tissues, using different animal models as well as human samples. In the context of regeneration, confocal Brillouin microscopy has recently been used to study spinal cord regeneration in the zebrafish (Schlüßler et al., 2018) and bone regeneration in rabbits (Akilbekova et al., 2018). Whereas atomic force microscopy (AFM)-based indentation measurements, the current gold standard in mechanobiology (Krieg et al., 2019), has been more extensively used in the field (Harn et al., 2021; Möllmert et al., 2020; Moyle et al., 2020), albeit with the limitation of measuring processed tissues from *ex vivo* explants. In urodeles, the only study on tissue mechanics was performed *in vitro*, demonstrating that wound closure of skin explants strongly relies on the substrate biomechanical properties (Huang et al., 2015). Hence, further *in vivo* studies need to be carried out to better understand the influence of mechanics on regeneration.

In this work, we measure for the first time the viscoelastic properties of a tetrapod limb *in vivo*. Using a confocal Brillouin microscope, we measured the mechanical properties of the cartilaginous skeleton during development and regeneration of the axolotl (*Ambystoma mexicanum*). The cartilage is a prominent structure in the limb that is formed/regenerated by chondroprogenitors which produce a specialized extracellular matrix (ECM), giving the cartilage its characteristic mechanical properties (Kozhemyakina et al., 2015). Here, we provide evidence that disruption of cartilage integrity results in a sustained decrease of the Brillouin frequency shift values. In addition, we detected a continuous decrease in tissue compressibility during limb development and cartilage differentiation in the digit, revealed by increasing Brillouin frequency shift values. Furthermore, we quantitatively mapped the mechanical properties inside the developing limbs and digits of live animals and were able to identify several anatomical structures, such as different stages of cartilage condensation, interstitial space and epidermis. Finally, we followed regeneration *in vivo*, demonstrating changes in the mechanical properties of the cartilage at different time points after amputation. Specifically, we detected an initial decrease in Brillouin frequency shift values in early stages of regeneration, which later increased approaching the levels of intact digits.

This work constitutes the first step for the application of Brillouin microscopy as a mean to characterize the mechanical properties in the highly regenerative axolotl. Taking advantage of this animal model, future *in vivo* studies will certainly open new possibilities to further comprehend the complexity of the regenerative process.

## MATERIALS AND METHODS

### Animal husbandry and handling

Husbandry and experimental procedures were performed according to the Animal Ethics Committee of the State of Saxony, Germany. Axolotls were kept and bred in the axolotl facility of the Center for Regenerative Therapies Dresden (CRTD) of the Technische Universität Dresden (TUD). A full description of the husbandry conditions was recently published (Riquelme-Guzmán et al., 2021).

Animals were selected by their snout to tail length (ST), which is indicated individually in each experiment. Used transgenic reporter lines are indicated in Table I.

**Table I:**
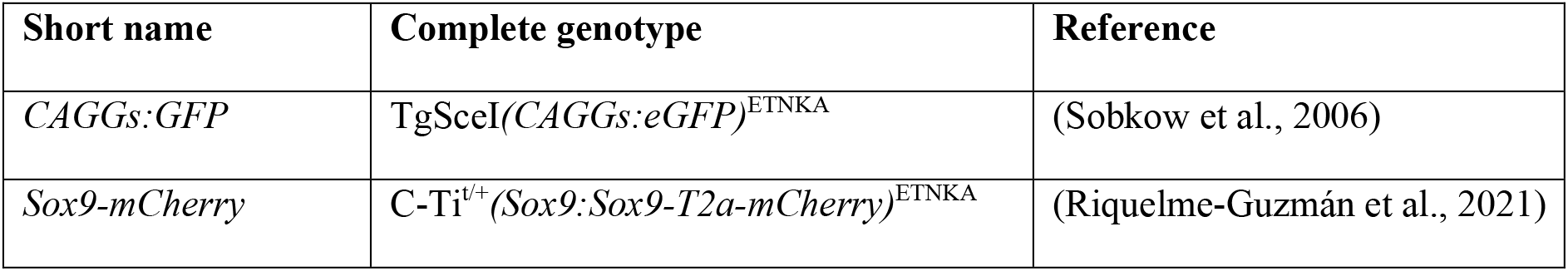
Transgenic reporter lines

The *CAGGs:GFP* line expresses *eGFP* driven by a modified cytomegalovirus-immediate early (CMV-IE) promoter, a strong and constitutively active promoter. Thus, eGFP is expressed in all cells (Sobkow et al., 2006). The *Sox9-mCherry* line has the mCherry protein fused to the endogenous SOX9 protein via a T2a self-cleaving peptide. Therefore, mCherry is present in all SOX9^+^ cells, which are particularly abundant in the cartilage (Riquelme-Guzmán et al., 2021).

Prior to all procedures, animals were anesthetized with 0.01% benzocaine solution diluted in tap water. After collection of tissues or when experiment was finished, animals were euthanized by exposing them to lethal anesthesia (0.1% benzocaine) for at least 20 minutes. Further experimental details are described in each case specifically.

### Confocal Brillouin microscopy and data analysis

Brillouin maps were acquired using a custom-built confocal Brillouin microscope previously described (Schlüßler et al., 2018). The setup is based on a two-stage VIPA interferometer. Illumination was achieved by a frequency-modulated diode laser with a wavelength of 780.24 nm. The laser frequency was stabilized to the D2 transition of rubidium 85. The setup has a CMOS camera (IDS UI-1492LE-M), which allows widefield images to be taken. Image acquisition was done with the custom-made software BrillouinAcquisition [Raimund Schlüßler and others (2017), BrillouinAcquisition version 0.2.2: C++ program for the acquisition of FOB microscopy data sets, available at https://github.com/BrillouinMicroscopy/BrillouinAcquisition]. Imaging was performed using a Zeiss Plan Neofluar 20x/0.5 objective, resulting in a spatial optical resolution of 1 μm in the lateral plane and 5 μm in the axial direction. The area of the imaged region and pixel size are indicated individually in each experiment.

Acquired data was analyzed using the custom-made software BrillouinEvaluation [Raimund Schlüßler and others (2016), BrillouinEvaluation version 1.5.3: Matlab program for the evaluation of Brillouin microscopy data sets, available at https://github.com/BrillouinMicroscopy/BrillouinEvaluation], and values in each map were exported to a CSV file. To analyze discrete areas within our Brillouin frequency shift maps, the custom-made software Impose was used [Paul Müller and others (2021), impose version 0.1.2: Graphical user interface for superimposing and quantifying data from different imaging modalities, available at https://github.com/GuckLab/impose].

The measured Brillouin frequency shift *ν_B_* can be expressed in terms of the longitudinal Modulus *M′*, refractive index *n* and density of the specimen, as well as the incident wavelength *λ*_0_ and scattering angle *θ* given by the setup: 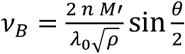. All measurements were performed in a backscattering configuration with *θ* = 180° and accordingly 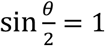. The longitudinal compressibility *κ_L_* may be expressed as the inverse of the longitudinal modulus 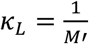. To facilitate the comparison with the results of other Brillouin microscopes, we normalized the measured Brillouin frequency shift *ν_B_* values with the frequency shift of water 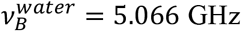. The dimensionless *Brillouin elastic contrast* 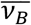 was previously introduced according to the following formula: 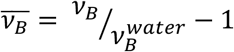 (Antonacci et al., 2020). 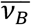 carries the same information as the Brillouin frequency shift *ν_B_*, but does not depend on the incident wavelength *λ*_0_.

### Imaging cartilage mechanical properties upon collagenase treatment *ex vivo*

For *ex vivo* measurements, we used the *CAGGs:GFP* reporter line (5-8 cm ST). One hand was collected and imaged immediately (no treatment) by placing it on a glass bottomed dish (ø: 50/40 mm, Willco Wells HBSB-5040) with a silica block laid on top to flatten it and improve light penetrance. To properly place the focus in the region of interest (proximal resting zone in distal phalanx of second digit), we used the fluorescence signal from our transgenic line and acquired a widefield image. After Brillouin image acquisition, hands were incubated with Liberase™ (0.35 mg/mL in 80% PBS, Sigma LIBTM-RO) for 30 minutes in a thermal mixer at 37 °C. Next, hands were rinsed briefly with 80% PBS and imaged again following the same steps mentioned before. A second incubation with the collagenase mix was performed for 15 minutes (45 minutes in total) prior to final imaging with the confocal Brillouin microscope. Generated maps correspond to a region of 148 x 80 μm, using a pixel size of 2 μm, averaging 2 acquisitions of 0.5 seconds *per* pixel.

### Imaging mechanical properties during digit development *in vivo*

To follow the development of the digit, we used the *Sox9-mCherry* line and selected larvae at developmental stages 45, 47, 49, 52 and 54, based on morphological parameters and mCherry expression pattern, as previously described (Nye et al., 2003; Purushothaman et al., 2019). Animals were anesthetized and later immobilized on top of glass bottomed dishes. They were mounted laterally to ensure proximity between the developing limb and the glass. Images were acquired with the Brillouin confocal microscope using a 5 μm step size and 0.5 seconds acquisition time. The size of the Brillouin map was not constant, as animals grow fast during early phases of development (Riquelme-Guzmán et al., 2021). Thus, the acquisition area was defined according to the size of the structure of interest.

After imaging, animals were euthanized and fixed with MEMFa (MOPS 0.1 M pH 7.4, EGTA 2 mM, MgSO_4_-7H_2_O 1 mM, 3.7% formaldehyde) overnight at 4 °C, washed with PBS and delipidated with a solution containing sodium dodecyl sulfate (SDS) 4% and boric acid 200 mM, pH 8.5, on a rocker at 37 °C for 20 minutes. Next, the embryos were washed with PBS and stained with Hoechst 33342 (1:2000 in PBS, Invitrogen H3570) for 1 hour, after which embryos were cleared by an overnight wash at room temperature with a refractive index matching solution (RIMS, EasyIndex RI 1.52). Cleared embryos were mounted in the same way as for Brillouin imaging and were imaged on an inverted Zeiss confocal laser scanning microscope LSM780 using a Plan-apochromat 10x/0.45 objective.

### Imaging mechanical properties during digit regeneration *in vivo*

For digit regeneration experiments, we used the *Sox9-mCherry* line (4 cm ST). Animals were anesthetized and imaged by placing them on top of glass bottomed dishes. The hand was positioned properly prior to be flattened and immobilized. A wet tissue with benzocaine was placed on top of animals during imaging to prevent them from drying. After measurements with the Brillouin confocal microscope, animals were imaged *in vivo* with an inverted Zeiss confocal laser scanning microscope LSM780 (Plan-apochromat 20x/0.8) to reveal the mCherry^+^ cartilage. Imaging with both microscopes was performed before, at 15- and 30-days post amputation (0, 15 and 30 dpa, respectively).

Amputation was performed at the joint of the distal phalanx in the second digit. After surgical procedure, the animals were covered with a wet tissue (with benzocaine) and allowed to recover for 10 minutes, before transferring them back to holding water.

Generated maps correspond to a 200 x 80 μm region, using a 2.5 μm step size, averaging 2 acquisitions of 0.5 seconds *per* pixel. To correlate our Brillouin maps with regeneration progress, we used *Sox9-mCherry* siblings that were amputated within the same day as the ones used for Brillouin imaging. Animals were imaged with an Olympus SZX10 stereoscope, using the Olympus cellSens Entry software. We measured the length of the digits from the joint to their distal-most tip and normalized their length with respect to day 0 (both for regenerating as well as undamaged digits).

### Brillouin data representation and statistical analysis

All Brillouin maps were generated by graphing the exported data as heat maps with GraphPad Prism Software. All other graphs and statistical analysis were generated with the same software. Specific statistical analysis is detailed in the corresponding figure legends, where we considered a *p*-value below 0.05 as significant.

To calculate the average Brillouin frequency shift within a defined region of interest, the values were extracted from the Brillouin maps, averaged *per* animal and plotted for multiple comparisons.

To generate the linear signal distribution graphs during development (Fig. 3F), five consecutive transversal lines were drawn in each Brillouin frequency shift map acquired for developing limb buds or digits. In all cases, drawn lines had a width of 1 μm and were separated by 5 μm between them. Brillouin frequency shift values were extracted from each line and averaged, generating one line *per* sample. Considering the differing sizes of limb buds and digits during development, structure size was normalized, with its center set as point zero and both edges as ±100%. Edges were defined by Brillouin frequency shift values below 5.1 (equivalent to benzocaine solution). Values from one representative individual *per* developmental stage were plotted together for qualitative comparison of changes occurring during development.

Images obtained with the confocal microscope or the widefield set-up of the Brillouin microscope were processed with the open source software Fiji (https://imagej.net/software/fiji/) (Schindelin et al., 2012). The images from developing limbs/digits correspond to a maximum intensity Z-stack projection. The Hoechst-derived nuclear signal was used to generate a mask to show the outline of the developing limbs/hands, which is displayed together with the mCherry signal, revealing *Sox9* expressing cells. The images of regenerating digits correspond to a maximum intensity Z-stack projection of four optical planes (1 μm interval), which correlates to the thickness of the section measured with the Brillouin confocal microscope (Schlüßler et al., 2018).

## RESULTS

### Confocal Brillouin microscopy maps cartilage architecture, identifying differences between chondrocytes and extracellular matrix

*In vivo* experiments in the axolotl are largely possible thanks to its transparency, particularly during juvenile stages, allowing the study of regeneration dynamics by using a diverse array of microscopy techniques. To evaluate how efficiently the mechanical properties in axolotl tissues can be probed, we tested the cartilaginous phalanxes using a confocal Brillouin microscope (Fig. 1A). Structurally, the cartilage is a tissue devoid of blood vessels and nerves, formed by chondrocytes embedded in a dense ECM, which contains in its majority collagen and, to a lesser extent, proteoglycans (Fox et al., 2009). In juvenile axolotls, appendicular skeleton is cartilaginous (Riquelme-Guzmán et al., 2021) and this cartilage accounts for up to 50% of the exposed surface upon amputation (Hutchison et al., 2007).

**Figure 1:**
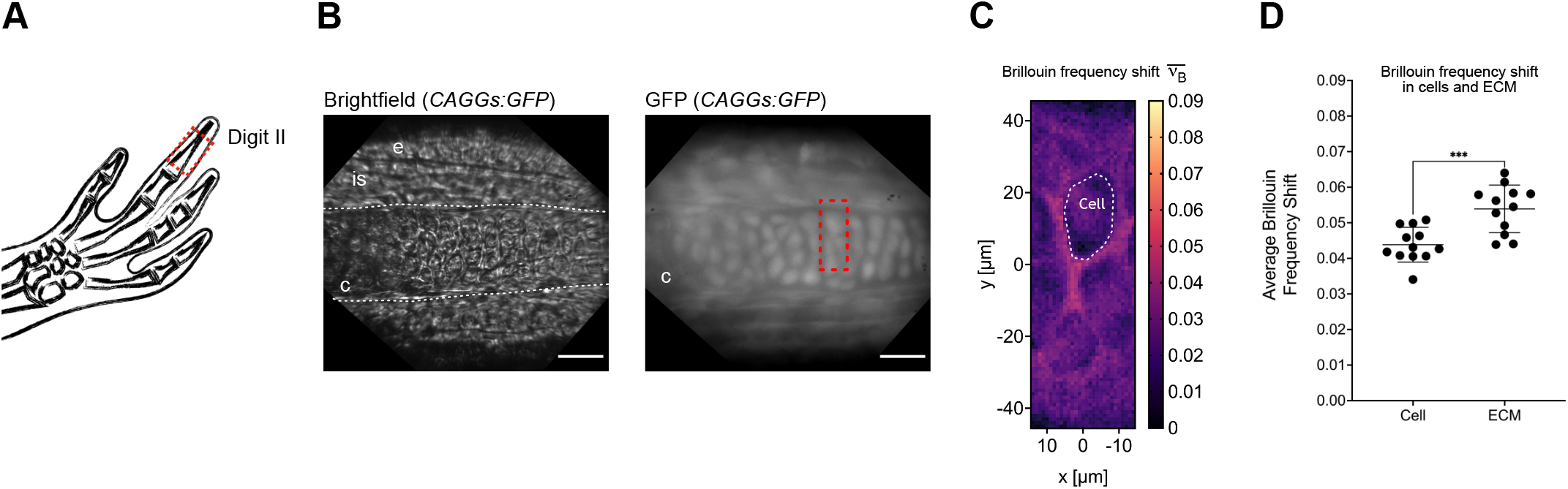
Confocal Brillouin microscopy maps cartilage architecture, identifying differences between chondrocytes and extracellular matrix. (A) Schematic representation of the distal phalanx in digit II. The red rectangle indicates the imaged area, shown in B. (B) Left: Brightfield image of probed region. Dashed lines show the position of the phalanx. Right: Widefield fluorescence image using *CAGGs:GFP* reporter line. All cells were eGFP^+^. Red rectangle show region probed with the Brillouin microscope shown in (C). c: cartilage, e: epidermis, is: interstitial space. Scale bar: 50 μm. (C) Brillouin elastic contrast 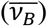 maps for the region shown in (B). The Brillouin elastic contract (normalized Brillouin frequency shift) is indicative of the elastic properties. Dashed oval show segmented cell as an example of quantifications shown in (D). (D) Quantification of average Brillouin frequency shift inside cells (chondrocytes) and the surrounding ECM (n = 12, *** p < 0.001, paired t-test).

To facilitate imaging the cartilage, we used *CAGGs:GFP* transgenic animals. This line expresses *eGFP* constitutively in all cells, allowing us to morphologically distinguish different tissues and structures (Fig. 1B). Particularly, the phalanx is an elongated structure with oval-shaped cells distributed along it (i.e., chondrocytes). Using 4 cm animals and a widefield set up, we could properly position the focus in a region of interest (ROI, Fig. 1B, red rectangle) where the mechanical properties were assessed using the confocal Brillouin microscope. As shown in Fig. 1C, we successfully mapped the Brillouin elastic contrast 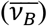 of a region inside the digit cartilage. The Brillouin elastic contrast (i.e., normalized Brillouin frequency shift) is indicative of the elastic properties of the sample. In the generated maps, we identified chondrocytes by morphology, with lower values than the surrounding ECM. In agreement with former reports (Bevilacqua et al., 2019), our results suggest that the collagen-rich matrix is less compressible as compared to chondrocytes. Moreover, by creating a mask to independently quantify the Brillouin elastic contrast of cells and the ECM, we found that indeed the ECM had significantly higher values than the chondrocytes embedded within (Fig. 1D).

Altogether, we were able to successfully probe the mechanical properties of the cartilage in axolotl digits, providing sufficiently high resolution to distinguish differences between intracellular and extracellular environments.

### Confocal Brillouin microscopy is sensitive to changes in cartilage architecture

Having demonstrated that the cartilage architecture can be mapped with confocal Brillouin microscopy, we wondered if changes in tissue structure could be assessed with this technique as well. We thus developed a protocol to follow the loss of tissue integrity via enzymatic treatment. As most of the ECM in cartilage is made of collagen, we treated the tissue with a protease mix enriched in collagenases (Liberase™). We collected hands from 5-8 cm *CAGGs:GFP* animals and incubated them with the collagenase mix for 30 and 45 minutes at 37 °C. For imaging, we always selected the proximal resting zone (RZ) of the distal phalanx in the second digit (Fig. 2A). The RZ corresponds to a region in which rounded quiescent chondrocytes are found at both ends of the skeletal element. As observed in the fluorescent microscopy images, treatment with collagenase did not largely affect the organization of chondrocytes at 30 minutes, but at 45 minutes a higher disruption of the tissue was observed, together with a reduction in cell density (Fig. 2B). Noteworthy, positioning the sample in the RZ was still possible, as the overall structure of the phalanx remained distinguishable.

**Figure 2:**
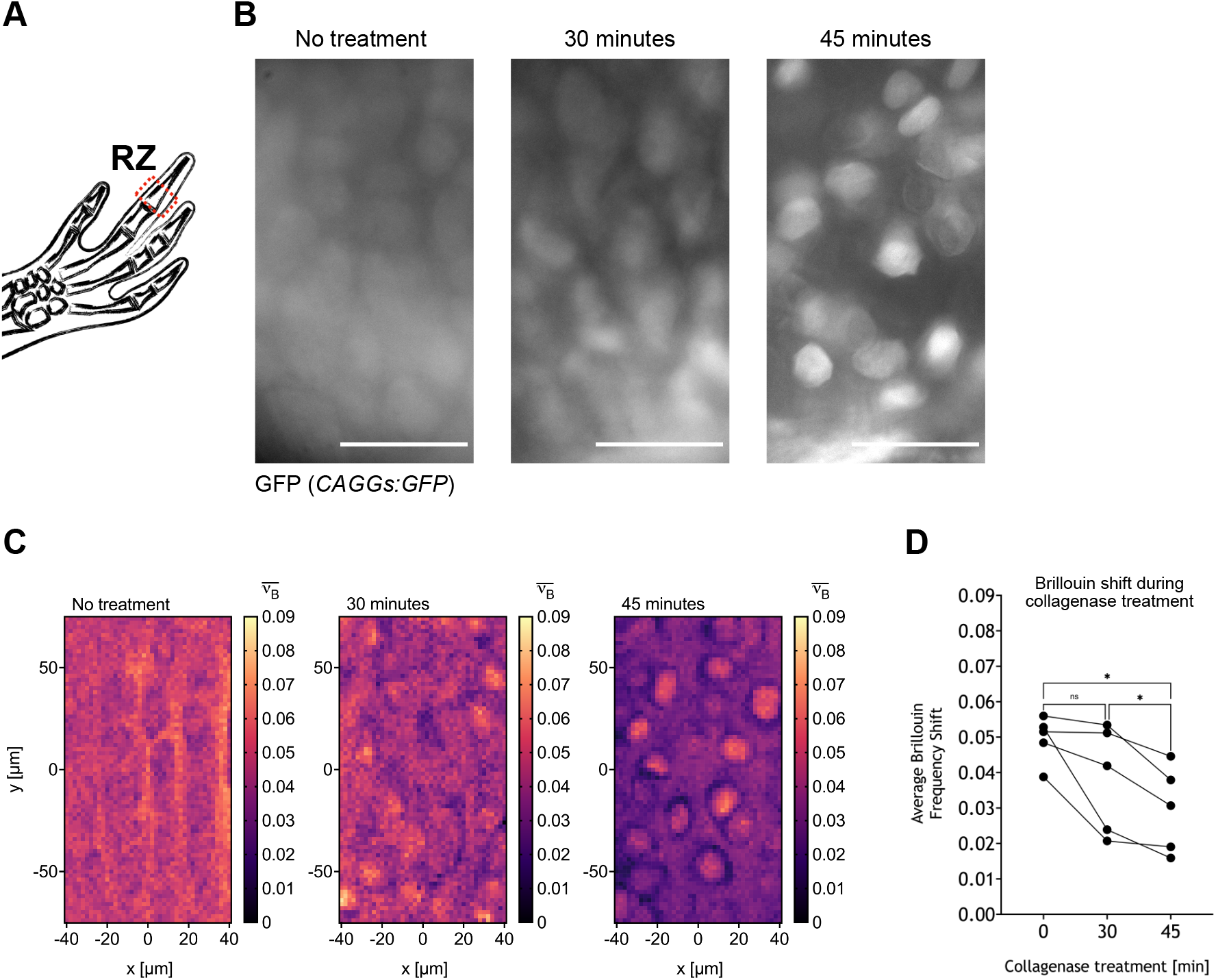
Loss of tissue integrity assessment with confocal Brillouin microscopy. (A) Schematic representation of proximal resting zone (RZ) of the distal phalanx in digit II. (B) Widefield fluorescence images of region in *CAGGs:GFP* hands treated with collagenase at different times, which were probed using confocal Brillouin microscopy. Scale bar: 50 μm. (C) Brillouin elastic contrast 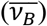 maps for the regions shown in (B). Maps reveal a time-dependent loss of tissue integrity upon collagenase treatment. (D) Average 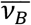 values of maps generated from *CAGGs:GFP* hands treated with collagenase (n = 5, * p < 0.05, one-way ANOVA with Tukey’s multiple comparisons test).

Using the confocal Brillouin microscope, we imaged a 148 x 80 μm region with a pixel size of 2 μm in the RZ (Fig. 2C). The Brillouin elastic contrast map in the RZ for untreated samples was similar to the one in Fig. 1C, with higher values in the ECM compared to the chondrocytes. Upon treatment for 30 minutes, a decrease in the Brillouin frequency shift was evident throughout the sample, particularly in the ECM. At 45 minutes, the values in the ECM were further decreased, allowing the cells to be easily visible in the map. Interestingly, the chondrocytes seemed to round up after the protease treatment (Fig. 2B), accompanied by an increase of their Brillouin frequency shift values. A similar phenomenon has been shown to occur in mesenchymal stem cells, as they stiffen immediately after detachment (Maloney et al., 2010).

To quantitatively compare the changes in the mechanical properties during the enzymatic treatment, we quantified the Brillouin frequency shift in the whole region by averaging all pixels in the map (Fig. 2D). Congruent with our qualitative observations, we detected a significant reduction in the Brillouin frequency shift values after 45 minutes of enzymatic treatment as compared to time 0 and 30 minutes. Overall, using the Brillouin microscope, we were able to spatially map the changes in the tissue architecture and mechanical properties of axolotl cartilage *ex vivo* upon protease treatment, as well as quantifying the increase in tissue compressibility during its loss of integrity.

### The Brillouin frequency shift increases progressively during digit development

To assess the physiological mechanical changes in the axolotl, we imaged the formation of the digit at different developmental stages *in vivo*. Using the *Sox9-mCherry* transgenic line, we have shown mCherry+ cells to be found in all the cartilage (Riquelme-Guzmán et al., 2021).

We collected embryos at stages 45, 47, 49, 52 and 54 based on previous work (Nye et al., 2003; Purushothaman et al., 2019). Embryos were imaged with the confocal Brillouin microscope and subsequently collected, fixed and cleared in order to perform confocal imaging of the mCherry^+^ region. At stage 45, no zeugopod is formed, and only a condensation of the humerus was observed (Fig. 3A). The zeugopod was observed at stage 47, when condensation of the radius and ulna became evident (Fig. 3B). From stage 49 onwards, condensation of the autopod skeletal elements were detected. Digits I and II were observed at stage 49 (Fig. 3C), followed by digit III at stage 52 (Fig. 3D) and digit IV at stage 54 (Fig. 3E). We measured with the confocal Brillouin microscope (pixel size 5 μm) in the distal-most part of the limb bud (stages 45 and 47) and, when the digits formed, in the distal end of digit II (Fig. 3A-E, red squares) assessing the change in mechanical properties during development.

**Figure 3:**
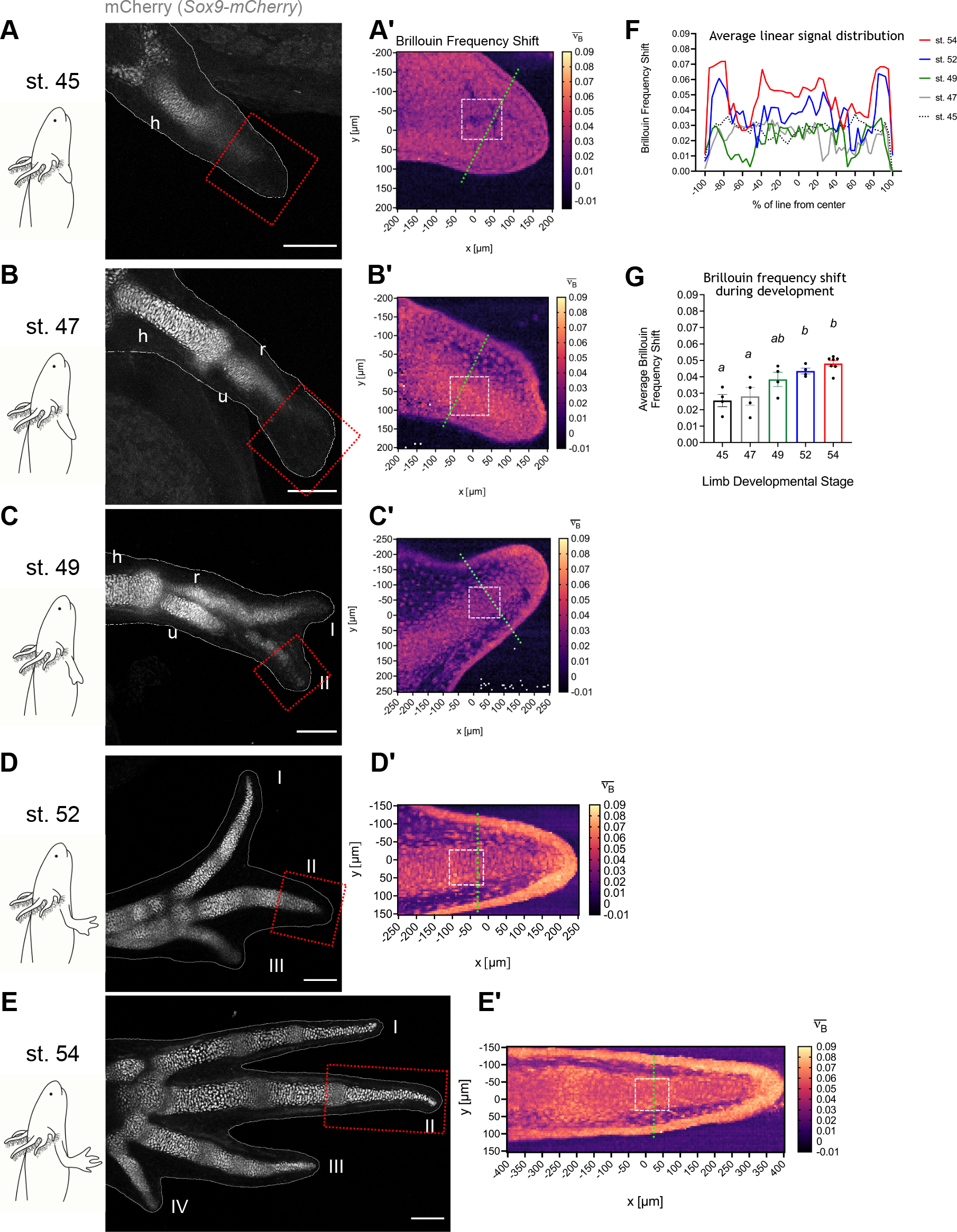
The Brillouin frequency shift increases progressively during digit development. (A-E) Left: schematic representation of limb development stages. Right: Confocal imaging of cleared *Sox9-mCherry* transgenic animals after Brillouin acquisition for each developmental stage. White dashed lines show limbs outline derived from nuclear staining with Hoechst. The red square represents the region probed with the Brillouin microscope. h: humerus, u: ulna, r: radius, I-IV: digit I – IV. Scale bar: 200 μm. (A’-E’) Normalized Brillouin frequency shift 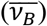 maps for regions shown in (A-E). Green dashed lines represent zones quantified in (F). White squares represent regions quantified in (G). Different structures in the digit were identified by their differences in the Brillouin frequency shift values. (F) Average 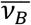 linear distribution along 5 transversal lines across the limb bud or digit at different developmental stages (n = 1 representative animal/developmental stage). (G) Average 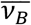 of regions shown in white squares in (A’-E’) (n ≥ 4 animals/developmental stage, each letter represents statistically equivalent values, one-way ANOVA with Tukey’s multiple comparisons test).

At stage 45, the limb bud is a semi-homogeneous structure with practically no perceivable differences revealed by the Brillouin elastic contrast map (Fig. 3A’), although a thin layer at the edge of the limb bud could be distinguished, likely corresponding to its epithelium. At stage 47, the map showed a slight increase in Brillouin frequency shift values in the middle region of the developing limb (Fig. 3B’). This increase could point towards early steps of cartilage condensation, which was not detected with the reporter of SOX9 expression (mCherry^+^ cells). In stages 49, 52 and 54, the digit II was morphologically identified at different developmental stages using mCherry expression. Our results show that the Brillouin frequency shift values progressively increased in the phalanx, as well as in the epidermis, during digit development (Fig. 3C’-E’), suggesting a decrease in tissue compressibility. Moreover, the Brillouin elastic contrast maps allowed us to recognize other structures such as the interstitial space, with likely some cells scattered throughout the tissue. At stage 54, we could also distinguish between the phalanx and joint. Additionally, we clearly identified the epidermis in all stages, together with a clear thickening from stages 49 to 54.

Interestingly, we observed in developing digits that Brillouin frequency shift values were higher in chondrocytes when compared to their surrounding ECM (Fig. 3C’-E’). This contrasts with what we observed in larger animals, where the ECM presented higher values than the chondrocytes embedded within (Fig. 1 and 2). This might be explained by development-associated changes in the cartilage matrix composition and deposition, which leads to progressively decreasing ECM compressibility (reviewed in Fowler & Larsson, 2020).

To further evaluate the mechanical differences within the limb bud and digit, we quantified the signal along several transversal lines (Fig. 3A’-E’, green dashed line) for every generated Brillouin elastic contrast map (n = 5 lines/animal). From these lines, we generated an average for each developmental stage (Fig. 3F). We observed a progressive increase in the average signal distribution with development. Moreover, this unidimensional data representation reveals the structures identified in our maps from stage 49 onwards (i.e., phalanx, interstitial space and epidermis) by their differences in the Brillouin frequency shift.

Finally, to quantitatively assess the differences in the Brillouin frequency shift during cartilage development, the signal within a 100 x 100 μm square region was averaged *per* animal (Fig. 3A’-E’, white dashed square). The regions selected correspond to mCherry^+^ tissue of digit II or to the distal end of the limb bud in stages 45 and 47 as controls (Fig. 3G). As expected, a progressive increase in the Brillouin frequency shift was observed in each developmental stage, suggesting a gradual decrease of cartilage compressibility. This observation is likely correlated with the increased ECM deposition and cell density that naturally occur during cartilage development (Fowler and Larsson, 2020).

Taken together, these results demonstrate the power of the confocal Brillouin microscope to probe mechanical properties in the axolotl *in vivo*. We were able to characterize the development of complex structures such as the digit, and to detect changes in tissue mechanics that were not measurable by the mere expression of transgenes.

### Brillouin frequency shift increases as cartilage condensates during digit regeneration

Considering our results hitherto, we next assessed if the confocal Brillouin microscope could also resolve mechanical changes occurring during regeneration. Thus, we acquired Brillouin frequency shift maps of the cartilage during digit regeneration *in vivo*. For this, we amputated at the joint of the distal phalanx from the second digit in 4 cm axolotls (Fig. 4A). Using the *Sox9-mCherry* line, we were able to properly position the sample and thus image the condensation of the regenerating cartilage. After every measurement with the confocal Brillouin microscope, we performed live confocal fluorescence imaging in order to observe the morphology of the regenerating digit at the same optical plane (Fig. 4B). Before amputation (0 dpa), we observed the RZ of the mature phalanx which was located distal to the amputation plane. This zone corresponded to well-organized rounded chondrocytes. At 15 dpa, condensation had already started, evidenced by characteristic tightly packed mCherry^+^ cells, forming an elongated structure. Finally, at 30 dpa, regeneration was in its later stages. A clearly defined digit was observed and the mCherry^+^ cells presented a morphology resembling that of rounded chondrocytes from a mature digit (i.e., 0 dpa).

**Figure 4:**
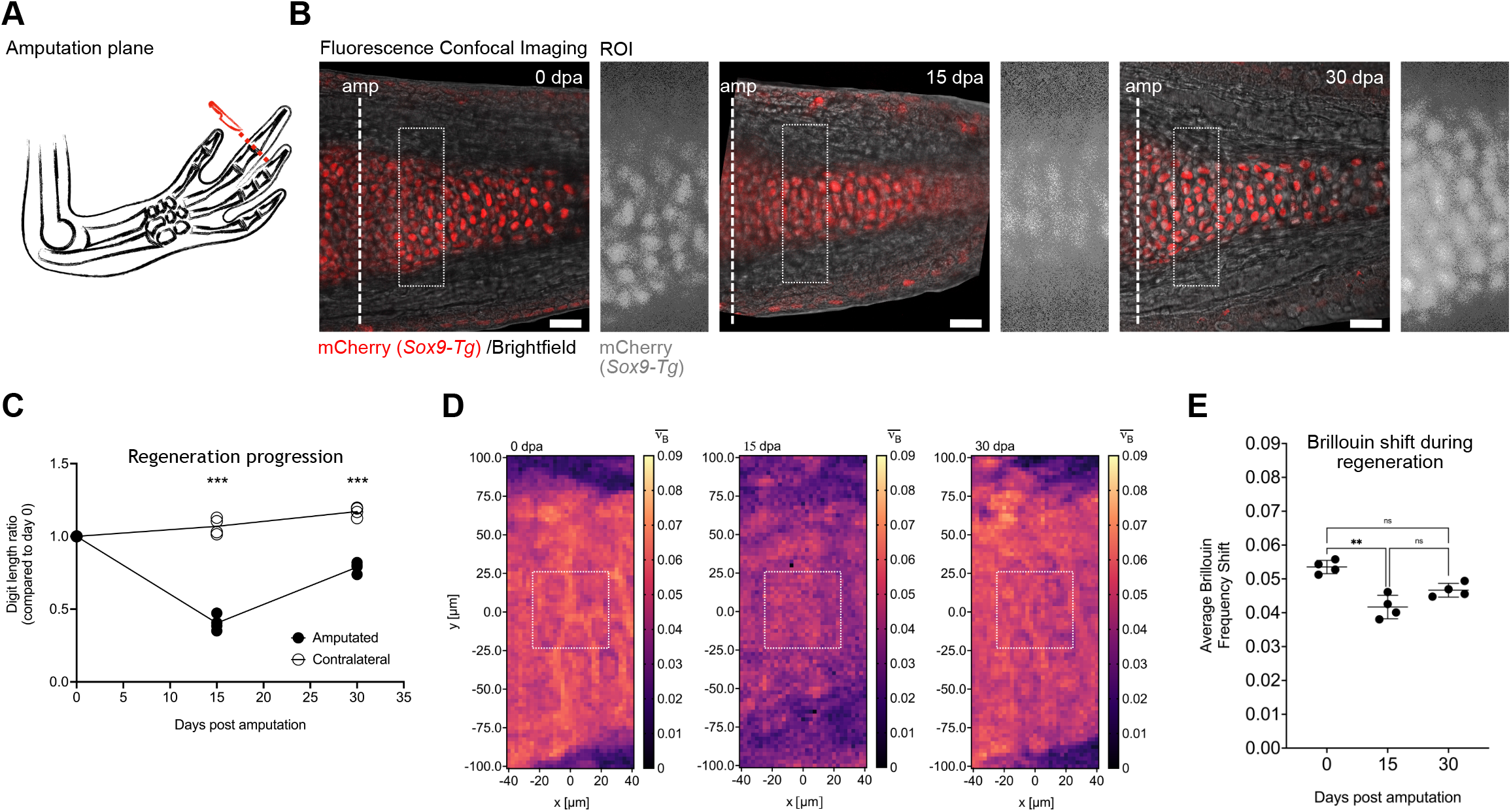
Brillouin frequency shift increases as cartilage condensates during digit regeneration. (A) Schematic representation of amputation plane at the joint of the distal phalanx. (B) Confocal live imaging of digit II from *Sox9-mCherry* axolotl before amputation (0 dpa), at 15- and 30-days post amputation (dpa). Images represent a composite between brightfield and maximum intensity Z-stack projection of 4 optical slices for the mCherry signal. Dashed line at 0 dpa shows the amputation plane (amp). Dashed rectangles show region of interest (ROI) which was probed with the confocal Brillouin microscope. A widefield fluorescence image taken with Brillouin set-up of the ROI is shown to the right of each timepoint. Scale bar: 50 μm. (C) Regeneration progression as shown by the digit length ratio at each timepoint. Ratio was calculated comparing each digit length with respect to day 0. Line is connecting mean values (n = 4, *** p < 0.001, amputated *vs* contralateral, two-way ANOVA with Bonferroni’s multiple comparisons test). (D) Brillouin elastic contrast 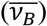 maps for ROIs shown in (B). White squares represent the region quantified in (E). Changes in 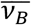 values were observed inside the cartilage at different timepoints. (E) Average 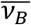 in cartilage during regeneration (n = 4, ** p < 0.01, one-way ANOVA with Tukey’s multiple comparisons test).

To quantify the general regeneration progress, we measured the length of the digit at these three time points and compared them to the contralateral one, i.e., the undamaged digit II of the opposite hand (Fig. 4C). A clear increase in the digit length was observed, which correlated with the progression of the regenerative process. Of note, even at 30 dpa, the length of the regenerated digit was significantly shorter than the contralateral one, indicating that the digit was not fully regenerated yet.

To compare the mechanical properties of the cartilage at these three different time points during regeneration, we measured the Brillouin frequency shift in the mCherry^+^ regions distal to the amputation plane (Fig. 4B, dashed rectangle). Similar to the abovementioned experiments, we were able to properly position the focus of the confocal Brillouin microscope using the mCherry signal and thus generate a Brillouin elastic contrast map covering mostly the cartilaginous tissue. The image corresponding to the acquired ROI was also used to identify the correct imaging plane in the confocal fluorescence microscope.

By creating a 200 x 80 μm map with a 2.5 μm pixel size, we were able to properly distinguish the differences of the mechanical properties of the cartilage during regeneration (Fig. 4D). At 0 dpa, the ECM had a higher Brillouin frequency shift than the chondrocytes within, and the interstitial tissue had lower values than the phalanx. These observations were in agreement with our observations in intact mature digits in previous sections. At 15 dpa, the Brillouin frequency shift was importantly reduced, with no clear distinction between the ECM and the chondrocytes, albeit the cartilage could still be identified by presenting slightly higher values than the interstitial tissue. This observation might be due to the stage of differentiation, as the condensation of the cartilage at this time point was in its earliest steps. Finally, at 30 dpa we observed a re-establishment of the mechanical properties in the phalanx, with a clear increase in the Brillouin frequency shift in the cartilage, evidenced by the color scale in the map. This map showed a similar trend to the uninjured digit, in which the ECM presented higher values than the cells embedded within.

To quantitatively assess the differences in the mechanical properties, we selected a 25 x 25 μm region in the center of each phalanx (Fig 4D, white squares) and averaged the values of all pixels within one region *per* animal (Fig. 4E). Congruent with our observations, we quantified a significant reduction in the Brillouin frequency shift at 15 dpa compared to day 0, which suggests that the cartilage might be more compressible during the early stages of condensation in the regenerating digit. When we analyzed the values at 30 dpa, we did not detect any significant difference with respect to either 0 or 15 dpa, suggesting that the tissue was in a transitioning stage, not yet reaching its final maturation. Thus, we could speculate that at later time points, the average Brillouin frequency shift would reach equivalent values to the ones acquired in uninjured tissues. These observations contrast with the interpretation of our images obtained with confocal fluorescence imaging, that suggested a re-establishment of the digit morphology after 30 days of regeneration (Fig. 4B, 30 dpa), thus revealing the strong capacity of Brillouin imaging for the detection of changes in viscoelastic tissue properties, which may be overlooked with traditional imaging techniques.

Collectively, our results provide novel evidence for measurement of viscoelastic properties in the axolotl *in vivo*. We demonstrated how the tissue architecture can be described through mechanical parameters such as compressibility, represented by the Brillouin elastic contrast. Additionally, we have shown that the loss of tissue integrity can be detected through a change in tissue architecture, as well as an overall decreased Brillouin frequency shift. We have also demonstrated for the first time that the tissues display increasingly reduced compressibility during the development of limbs and digits *in vivo*. Furthermore, using the confocal Brillouin microscope, we were able to distinguish differences in the mechanical properties of the cartilage during digit regeneration at different timepoints.

## DISCUSSION

In recent years, confocal Brillouin microscopy has emerged as a powerful label-free and contact-free tool to probe mechanical properties of biological samples (reviewed in Prevedel et al., 2019). Measurements of diverse systems have been carried out *in vitro* and *ex vivo*; however, the adaptation of confocal Brillouin microscopy for *in vivo* animal models has only been performed using zebrafish larvae to study spinal cord repair and growth (Schlüßler et al., 2018), the material properties in the notochord (Bevilacqua et al., 2019) and retina development (Amini et al., 2021). In this report, we present for the first time an assessment of the *in vivo* mechanical properties of the axolotl, both during development and regeneration. Thereby, we add to the existing literature by providing yet another application of Brillouin microscopy to the field of tissue mechanics.

The composition and structure of the extracellular matrix (ECM) entails tissue specific biochemical and biomechanical properties that play key roles in homeostasis and pathologies, such as cancer and fibrotic disorders (reviewed in Young *et al*, 2016). In addition, due to the pronounced contribution of the ECM to mechanical tissue properties, the ECM and its constituents have been targeted in various experiments employing Brillouin light scattering (Cusack and Lees, 1984; Cusack and Miller, 1979; Harley et al., 1977; Palombo et al., 2014). This also includes studies on the ECM-rich cartilage tissue, in which an important reduction of the Brillouin frequency shift was reported in porcine articular cartilage treated with trypsin, due to degradation of proteoglycans (Wu et al., 2019), the second largest structural component of the ECM (Fox et al., 2009). Moreover, distinctive Brillouin frequency shift values have been reported for different parts of the human femoral head, including the measurement of trabecular bone, subchondral bone and articular cartilage. These measurements revealed a prominent mechanical heterogeneity throughout all these regions (Cardinali et al., 2019).

In the present study, we mapped the Brillouin frequency shift in the resting zone of axolotl phalanges, which allowed us to identify distinctive mechanical properties of the cartilage ECM and embedded chondrocytes. In intact mature digits, the ECM displayed significantly higher Brillouin frequency shift values as compared to the chondrocytes. A similar mechanical heterogeneity was observed in zebrafish larvae where ECM-rich regions displayed higher values than neighboring cells (Bevilacqua et al., 2019). During development, we observed gradually increasing Brillouin frequency shift values from limb developmental stage 45 to 54. This suggests a progressive decrease in tissue compressibility as the cartilage matures, which may be explained by changes in ECM composition and structure (Fowler and Larsson, 2020). Changes of mechanical tissue properties during cartilage development have so far only been measured *ex vivo* (Gannon et al., 2015). By assessing tissue compressibility, we were able to observe this phenomenon for the first-time *in vivo*.

As collagen is the main component of the cartilage ECM, making up to 60% of its dry weight (Fox et al., 2009), we could perturb cartilage integrity and record a significant reduction of the Brillouin frequency shift after treatment with collagenase *ex vivo* in a time-dependent manner. This result is in agreement with the reduction of the Brillouin frequency shift reported in porcine articular cartilage after enzymatic degradation of proteoglycans (Wu et al., 2019).

Undoubtedly, tissue mechanics play a critical role in development, and given its parallels with regeneration (reviewed in Vining & Mooney, 2017), the role of mechanical tissue properties in the latter needs to be further explored. Accordingly, by successfully mapping cartilage viscoelasticity *in vivo* during development and regeneration, we detected important similarities between both processes. During development, the Brillouin frequency shift gradually increases as the animal grows, and during regeneration, after a transient decrease, we also detected an increase in the Brillouin frequency shift inside the cartilaginous structure. Such variations in viscoelastic cartilage properties indicate the dynamic environment, from a mechanical perspective, in which development and regeneration occur. Importantly, development and regeneration do not only involve cartilage, but also other tissues such as muscle, blood vessels and nerves. Therefore, it will be important to assess how mechanical tissue properties are contributing to the shape of the limb from a multi-tissue perspective, integrating the information that we have for individual tissues into one integral model.

Confocal fluorescence imaging is a powerful tool to identify biologically relevant molecules or structures at the subcellular and tissue-scale levels with high resolution (Elliott, 2020). However, here we have demonstrated that the expression of fluorescent proteins as an indicator of tissue architecture alone may not be sufficient. During our digit regeneration experiments, by acquiring Brillouin frequency shift maps, we show that the regenerative process was not completed mechanically by 30 days after amputation, which correlated with our regeneration progress evaluation. Contrastingly, the arrangement of SOX9 expressing cells indicated a mature-like morphology.

One major advantage of Brillouin microscopy is the possibility to access mechanical properties in a non-invasive manner, circumventing the introduction of potential artifacts caused by sample preparations prior to tissue mechanical measurements. When *in vivo* Brillouin images of the perispinal area of zebrafish larvae were compared with *ex vivo* tissue slices of the same region, differences in material properties of the sample were observed (Schlüßler et al., 2018), indicating that sample processing for *ex vivo* measurements might introduce diverse artifacts that could affect the results. In contrast, the values for Brillouin frequency shift in our cartilage samples *in vivo* and *ex vivo* did not change significantly (*ex vivo*: 5.317 ± 0.03 GHz; *in vivo*: 5.337 ± 0.01 GHz; *p* = 0.281 Student’s t-test). It is important to note that in our study the positioning, i.e. the direction of the measurement, was the same for the *in vivo* and *ex vivo* samples. This, and the large distance between the measured area (digit) and the sectioning plane for tissue collection (wrist) differ from the described sample preparation between *ex vivo* and *in vivo* tissues in (Schlüßler et al., 2018). Tissue-specific mechanical properties may also play a contributing role (Vining and Mooney, 2017). The major difference between *in vivo* and *ex vivo* Brillouin frequency shift measurements in zebrafish samples were detected in muscle (Schlüßler et al., 2018), whereas we focused on cartilage that is non-contractile and might be more resistant to rapid degradation than other tissue types. Consequently, further systematic *in vivo* and *ex vivo* measurements will be required to properly unravel how the mechanical properties of living tissues are changing not only during biologically relevant processes such as development and regeneration, but also upon certain preparatory procedures.

In regenerative biology, the axolotl is a key model organism to study limb regeneration. Its natural transparency during juvenile stages, together with the increasing availability of datasets, such as single-cell RNAseq (Gerber et al., 2018; Leigh et al., 2018) and the establishment of tools, such as the present work, provide new opportunities to investigate the role of tissue mechanics in regeneration. Additionally, a great body of research has been done on limb patterning and positional memory during regeneration (reviewed in Otsuki & Tanaka, 2021). The re-establishment of a complex structure such as the limb, requires the release of cells from their surrounding ECM, their de-differentiation, migration and proliferation, followed by cell differentiation and tissue patterning. These important changes are accompanied and influenced by forces produced by cell-cell and cell-ECM interactions (Boilly et al., 1990; Petreaca and Martins-Green, 2019). Hence, confocal Brillouin microscopy opens the possibility to quantify the changes of material properties that accompany these regenerative processes and ultimately contribute to shaping the new limb.

Altogether, our work expands the potential of Brillouin microscopy as a tool to probe the *in vivo* mechanical properties of biological samples to the axolotl. With these results, we aim to set the basis for utilizing this methodology to answer relevant questions in the field of salamander regeneration. In combination with a myriad of currently available tools, Brillouin microscopy will enable researchers to understand how mechanical tissue properties are shaping and influencing the regenerative process.

## ACKNOWLEDGMENTS

We would like to thank all members of the Sandoval-Guzmán Lab for continuous support and valuable feedback during the development of this work. We are also grateful to Anja Wagner, Beate Gruhl and Judith Konantz for their impeccable dedication to the axolotl care. TSG and CRG were supported by a DFG Research Grant (SA 3349/3-1). JG was supported by the DFG (SPP 2191 – Molecular mechanisms of functional phase separation, grant agreement number 419138906). CRG was supported by the DIGS-BB fellow award. SEJ was supported by the European Union’s Horizon 2020 research and innovation program under grant agreement No 101022810. This work was supported by the Light Microscopy Facility, a Core Facility of the CMCB Technology Platform at TU Dresden.

